# Development and characterization of a recombinant silk network for 3D culture of immortalized and fresh tumor-derived breast cancer cells

**DOI:** 10.1101/2022.12.22.521677

**Authors:** Caterina Collodet, Kelly Blust, Savvini Gkouma, Emmy Ståhl, Xinsong Chen, Johan Hartman, My Hedhammar

**Affiliations:** Division of Protein Technology, School of Biotechnology, KTH Royal Institute of Technology, SE-106 91 Stockholm, Sweden; Department of Oncology-Pathology, Karolinska Institutet, Stockholm, Sweden; Department of Clinical Pathology and Cancer Diagnostics, Karolinska University Hospital, Stockholm, Sweden

**Keywords:** breast cancer, SK-BR-3, MCF-7, MDA-MB-231, FN-silk network, 3D model, RNA-seq

## Abstract

Traditional cancer models rely on 2D cell cultures or 3D spheroids, which fail to recapitulate cell-extracellular matrix (ECM) interactions, a key element of tumor development. Existing hydrogel-based 3D alternatives lack mechanical support for cell growth and often suffer from low reproducibility. Here we report a novel strategy to make 3D models of breast cancer using a tissue-like, well-defined network environment based on recombinant spider silk, functionalized with a cell adhesion motif from fibronectin (FN-silk). With this approach, the canonical cancer cells SK-BR-3, MCF-7, and MDA-MB-231, maintain their characteristic expression of markers (*i*.*e*., ERα, HER2, and PGR) while developing distinct morphology. Transcriptomic analyses demonstrate how culture in the FN-silk networks modulates the biological processes of cell adhesion and migration while affecting physiological events involved in malignancy, such as inflammation, remodeling of the ECM, and resistance to anticancer drugs. Finally, we show that integration in FN-silk networks promotes the viability of cells obtained from the superficial scraping of patients’ breast tumors.

## 1. Introduction

Breast cancer is the most common cancer among women worldwide, excluding nonmelanoma skin cancer.^1^ Breast cancer is a heterogeneous disease composed of several subtypes, each with different morphological and clinical implications.^2^ To tailor therapies, patients are routinely classified by assessing the expression of three histological markers (*i*.*e*., ERα, HER2, and PR). In the past decades, several studies identified collections of genes that allow further patient stratification.^3^

Despite significant advances in the field, there is still a need to find new drugs. With many candidates failing in clinical trial phases,^4^ developing more reliable preclinical models is crucial. Traditionally, the first stages of preclinical research rely on cells grown on a flat surface, lacking the structure of tumors and tissues.^5^ Likely due to these oversimplified conditions, *in vitro* models often fail to recapitulate the *in vivo* response.^6,7^ Three-dimensional (3D) models have been suggested as a bridge between *in vitro* and *in vivo* models since they better mimic the complexity of the tumor microenvironment.^8,9^

3D models currently available can be divided into scaffold-free models, where cells are forced to self-aggregate forming tumor spheroids, and scaffold-based systems, with cells growing onto extracellular matrix (ECM)-mimetic biomaterials.^10^ Scaffold-free alternatives have been suggested to recapitulate the oxygen, nutrients, and pH gradients of solid human tumors^11^ but lack the cell-ECM interaction component. The alternative scaffold-based systems often rely on materials with ill-defined composition, having batch-to-batch variability and limited reproducibility.^12^ Additionally, it is often difficult to obtain good seeding efficiency due to cells inability to spread homogeneously through rigid structures.^13^ These drawbacks have driven the search for alternatives, one of which is the recombinantly produced spider silk protein 4RepCT functionalized to harbor the cell adhesion motif from fibronectin (FN),^14,15^ herein referred to as FN-silk. Cells can be added to an FN-silk solution before its assembly, which is then carried out under physiological conditions and leads to the formation of a network mimicking the fibrous part of the ECM with homogeneously integrated mammalian cells.^16^ Furthermore, FN-silk can be easily sterilized, is well tolerated *in vivo*, and of non-animal origin.^17^

In this study, we developed a new technique to generate floating networks of FN-silk containing breast cancer cells. This system extends our previous work in which various cell types, including endothelial cells, fibroblasts, keratinocytes, and pluripotent stem cells,^18^ were successfully cultured on FN-silk. For the current study, cell lines representative of the three main subtypes of breast cancer HER2-overexpression, luminal-like, and triple-negative, respectively SK-BR-3, MCF-7, and MDA-MB-231 were used to compare culture in FN-silk to traditional 2D culture. We assessed the proliferation rate and expression of the markers routinely used to classify breast tumors and investigated by RNA-sequencing (RNA-seq) how FN-silk modulates the transcriptional landscape of MCF-7 and MDA-MB-231. The bioinformatics analysis revealed that culture in FN-silk network affected the expression of genes related to pathophysiological cancer processes such as cell adhesion, migration, and inflammatory signaling. We also demonstrated that FN-silk networks are highly adaptable, allowing the growth of novel breast cancer cells, such as the clinically relevant PB and Wood, as well as cells obtained from fresh tumors.

## 2. Materials and methods

Full details are available in the Supplementary Information.

### FN-silk network formation

The FN-silk network was created using a silk protein functionalized with a motif from fibronectin, FN-silk (3 mg/ml in PBS, endotoxin level below 200 EU/ml), provided by Spiber Technologies AB. For each construct, 10.000 cells from a cell concentrate of 5.900 cells/μl were gently mixed with 8,3 μl FN-silk to form a 10 μl droplet. The droplet was placed on a polytetrafluoroethylene (PTFE) mold, previously anchored at the bottom of a 24-well plate. A foam was formed by rapidly pipetting air into the droplet. Constructs were stabilized by incubation for 20 min at 37°C. After, the FN-silk networks were transferred to a hydrophobic 96-well plate (Sarstedt, 83.3924.500) using a round-shaped stainless steel micro spoon (Sigma, Z648299). To facilitate the transfer, medium was added to the 24-well plate and the 96-well plate. After transferring, the medium was removed, and a 3D-printed cap was placed on the plate. The cap was then connected to a switched-off Vacusafe aspiration system (Integra). The Vacusafe was switched on for 1 min to allow for the creation of a pressure difference. Then the tubing part connected to the cap was quickly disconnected, allowing the pressure to release. This procedure was repeated twice. Fresh medium was added, and the constructs were kept in culture for seven days. The medium was changed every other day. 2D controls were created by plating 10.000 cells per well on 96-well plate TC treated (Thermo Fisher, 174929).

### Bioinformatic analysis

The gene ontology was performed using the over-representation analysis function for biological processes in Web-Gestalt.^19^ Additionally, a gene set enrichment analysis was done by submitting as a custom database the gene list of Nanostring nCounter® Breast Cancer 360. The transcription factor prediction was performed using i-*cis*Target^20^ and the upstream regulator function by QIAGEN Ingenuity Pathway Analysis.^21^ Differentially expressed genes (DEGs) comparing MCF-7 and MDA-MB-231 grown in FN-silk networks for a week with their counterpart kept in 2D were visualized as volcano plots using the ggplot2 package in R. The significance thresholds were set to the adjusted *p-* value < 0,05 and the log2 fold-change values ≥ |± 0,38|. The top 10 DEGs were highlighted (dark grey), and several important genes were marked (black).

### Accession numbers

The RNA-seq data that support the findings of this study are openly available in NCBI’s Gene Expression Omnibus at [https://www.ncbi.nlm.nih.gov/geo/query/acc.cgi?acc=GSE209570], reference number [GSE209570].

The code used to create the volcano plots is available on GitHub (https://github.com/kblust/vulcanoplot_breastcancer2Dvs3D).

## 3. Results

### The FN-silk networks offer an ECM-like environment for cells growth

Figure 1a describes the various steps required to form the FN-silk networks with breast cancer cells. An initial drop containing 10.000 breast cancer cells and FN-silk protein solution was placed onto a sheet of hydrophobic polytetrafluoroethylene (PTFE) in a culture well (step I). Air bubbles were introduced through rapid pipetting, allowing the creation of foam with multiple liquid-air interfaces in which FN-silk assembled into sheets around each air bubble (step II). The construct was placed in a cell incubator at 37°C for 20 minutes to stabilize the structure. Afterward, each foam was transferred to a well of a 96-well plate containing medium, using a micro spoon (step III). The medium was then removed (step IV), and, to promote the release of the air bubbles thereby yielding a network, a pressure difference was applied by placing a 3D-printed cap on top of the plate and connecting it to a standard medium aspiration tool for 2 minutes (step V). Finally, the medium was added, and cells were kept in culture for seven days within the FN-silk networks.

**Fig. 1.**
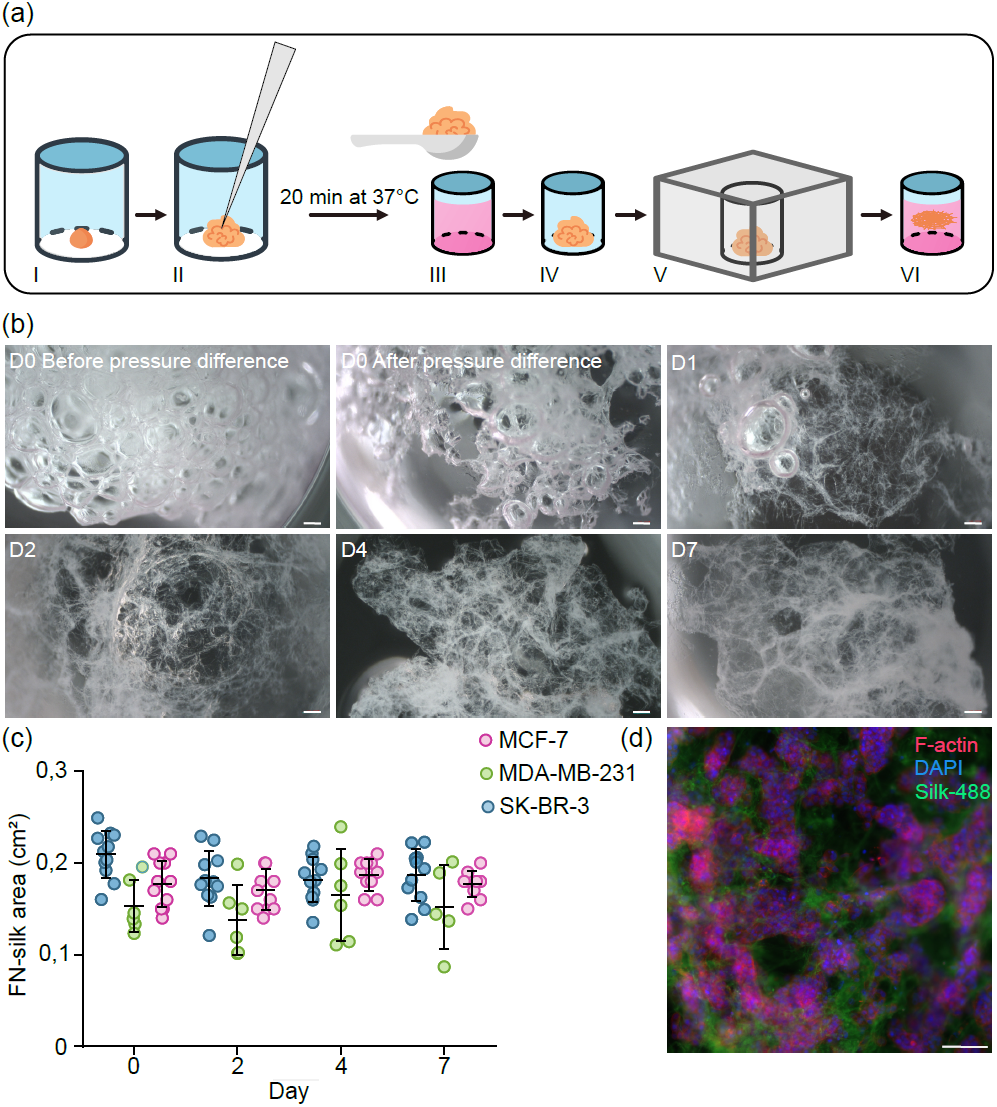
FN-silk network cancer model formation, network morphology, and area. (a) Cartoon depicting the protocol to form FN-silk network with cancer cells. Briefly, a solution comprising FN-silk and a cell suspension is deposited on a PTFE-coated well (I), self-assembly of FN-silk is promoted by air pipetting in the solution (II) after 20 min of stabilization at 37°C in the cell incubator, scaffolds are transferred with a micro spoon into the well of a 96-well plate (III), the medium is removed (IV), and a pressure difference is applied (V), the medium is added, and FN-silk networks are kept in culture for seven days (VI). (b) Macroscopic brightfield pictures of the FN-silk network on day 0 before and after pressure difference is applied and on days 1,2,4, and 7. Scale bar: 203 μm. (c) The area of FN-silk network constructs generated using the cell lines MCF-7 (pink), MDA-MB-231 (green), and SK-BR-3 (blue) is calculated by analyzing brightfield images taken at various time points (*i*.*e*., days 0, 2, 4, and 7). Average and standard deviations are shown for each time point; individual values are also indicated, (d) F-actin staining on MCF-7 cultured for 7 days on a scaffold made using FN-silk conjugated with 488-fluorophore. Overlay image of F-actin (red), nuclei (DAPI, blue), and FN-silk (green). Scale bar: 100 μm.

Brightfield pictures were taken at various time points to monitor how quickly the constructs would get to their final tissue-like appearance (fig. 1b). On the formation day (*i*.*e*., day 0), applying a pressure difference caused FN-silk to display an initial collapse of the air bubbles entrapped within the construct. After 24 hours, most of the air bubbles could no longer be detected and were completely gone 48 hours after formation, when the final ECM-like network was established. Pictures taken on days 4 and 7 confirmed that the FN-silk networks maintained their architecture over time.

Breast cancer is a heterogeneous disease with different subtypes associated with various prognoses and therapies. We chose to work with the three canonical human immortalized breast cancer cell lines SK-BR-3, MCF-7, and MDA-MB-231, representative respectively of the main subtypes HER2-overexpression, luminal-like, and triple-negative. To estimate the reproducibility among networks created with the three cell lines, we performed area measurements of the outer surfaces of the FN-silk networks. For each cell type, a set of ten scaffolds was generated, and stereomicroscopic pictures of each construct were taken on day 0, after formation, and on days 2,4, and 7. The measurements showed that for each cell type, the size of the scaffolds remained comparable over time. MDA-MB-231 scaffolds (area 0,13 cm^2^ ±0,05 cm^2^) appeared smaller than the ones generated with MCF-7 (0,18 cm^2^ ± 0,02 cm^2^) and SK-BR-3 (area 0,19 cm^2^ ± 0,02 cm^2^) cells (fig. 1c). An additional analysis done to estimate the area values independently of the cell line, revealed an average scaffold area of 0,17 cm^2^, corresponding to 60% of a 96-well plate well area (0,29 cm^2^) (Supplemental fig. 1a). The thickness of the networks was estimated to be between 150 to 200 μm by z-stack measurements done with fluorescent microscopy.

Finally, to examine the interactions established by the cells in the FN-silk network, we created scaffolds using a fluorescently-labelled FN-silk and let MCF-7 grow for seven days. Staining for cytoskeleton and nuclei of MCF-7 highlighted grape-like clusters of cells in close proximity to FN-silk, demonstrating how cells establish cell-cell and cell-matrix interactions (Fig. 1d).

### Breast cancer subtypes grow and maintain their key features in FN-silk networks

After verifying that breast cancer cells can establish cell-cell and cell-matrix contacts while growing within the FN-silk networks, we investigated the viability, marker expression, and morphology of cells.

Firstly, we confirmed a homogeneous distribution pattern by staining the cytoskeleton and nuclei of cells cultured in the scaffolds (Fig. 2a). We then performed the Alamar Blue assay at several time points (*i*.*e*., days 1, 4, and 7) to compare the metabolic activity of cells cultured in FN-silk networks or on tissue-culture treated plates. The data revealed that all cell lines could grow in FN-silk networks, with the highly invasive cell MDA-MB-231 having the highest metabolic rate. SK-BR-3 and MDA-MB-231 cultured in 2D showed a higher proliferation rate than their counterpart kept in FN-silk constructs, whereas MCF-7 had slightly higher metabolic activity in FN-silk networks after seven days in culture (Fig. 2b). Immunofluorescence staining for keratin 8 (KRT8), highlighted how MCF-7 formed mass-like clusters, in contrast to SK-BR-3 and MDA-MB-231, which appeared to spread without clumping together (Fig. 2c). This observation suggests a correlation between the higher metabolic rate in 3D and the ability of cells to develop complex morphologies in the ECM-like network.

**Fig. 2.**
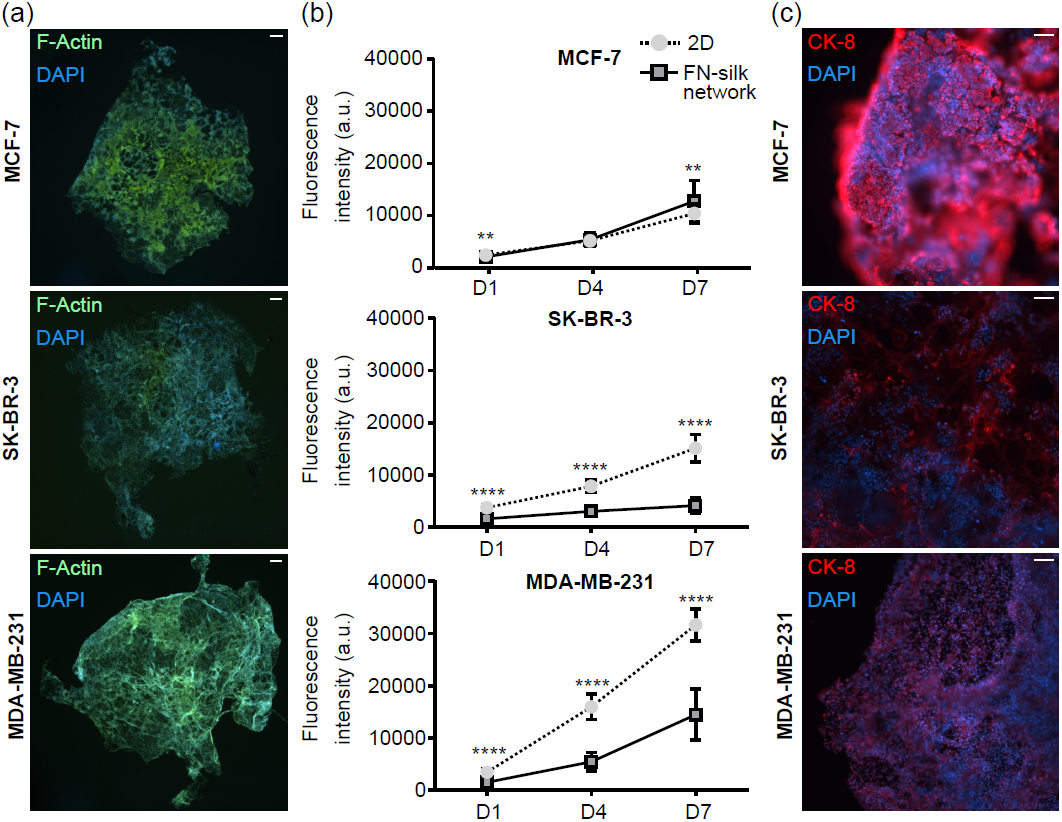
Breast cancer cells grow with a homogeneous distribution in FN-silk networks. (a) Overlay images of staining for actin filaments (F-Actin, green) and nuclei (DAPI, blue) were taken on MCF-7, SK-BR-3, and MDA-MB-231 cultured in FN-silk network for seven days. Scale bar: 100 μm. (b) Graphs indicating the metabolic activity of MCF-7, SK-BR-3, and MDA-MB-231 cells cultured in 2 dimensions (2D) or FN-silk network. The metabolic rate was monitored over time (days 1, 4, and 7) using Alamar blue. *\*\*P <* 0,01; *\*\*\*\*P <* 0,0001. (c) Overlay images of staining for cytokeratin 8 (CK-8, red) and nuclei (DAPI, blue) were taken on MCF-7, SK-BR-3, and MDA-MB-231 cultured in FN-silk network for seven days. Scale bar: 100 μm.

As it is crucial to maintain the features characterizing the heterogenicity of breast cancer subtypes, we next sought to assess the stability of the four classical markers estrogen receptor a *(ESR1*, ERa), progesterone receptor *(PGR*, PR), human epidermal growth factor receptor 2 *(ERBB2*, HER2) and, marker of proliferation Ki-67 *(MKI67*, Ki67) in our FN-silk models. The mRNA level measurement confirmed that cells maintain their characteristic features when cultured in FN-silk networks. For instance, we detected a high expression of *ERα* and *PGR* in the luminal-like MCF-7, and an abundant *ERBB2* in the HER2-overexpressing SK-BR-3, while these markers were absent in the triple-negative MDA-MB-231. The levels of *MKI67* emphasized the high proliferation of MDA-MB-231 and MCF-7 in contrast to a slower growth rate of SK-BR-3 (Fig. 3a).

**Fig. 3.**
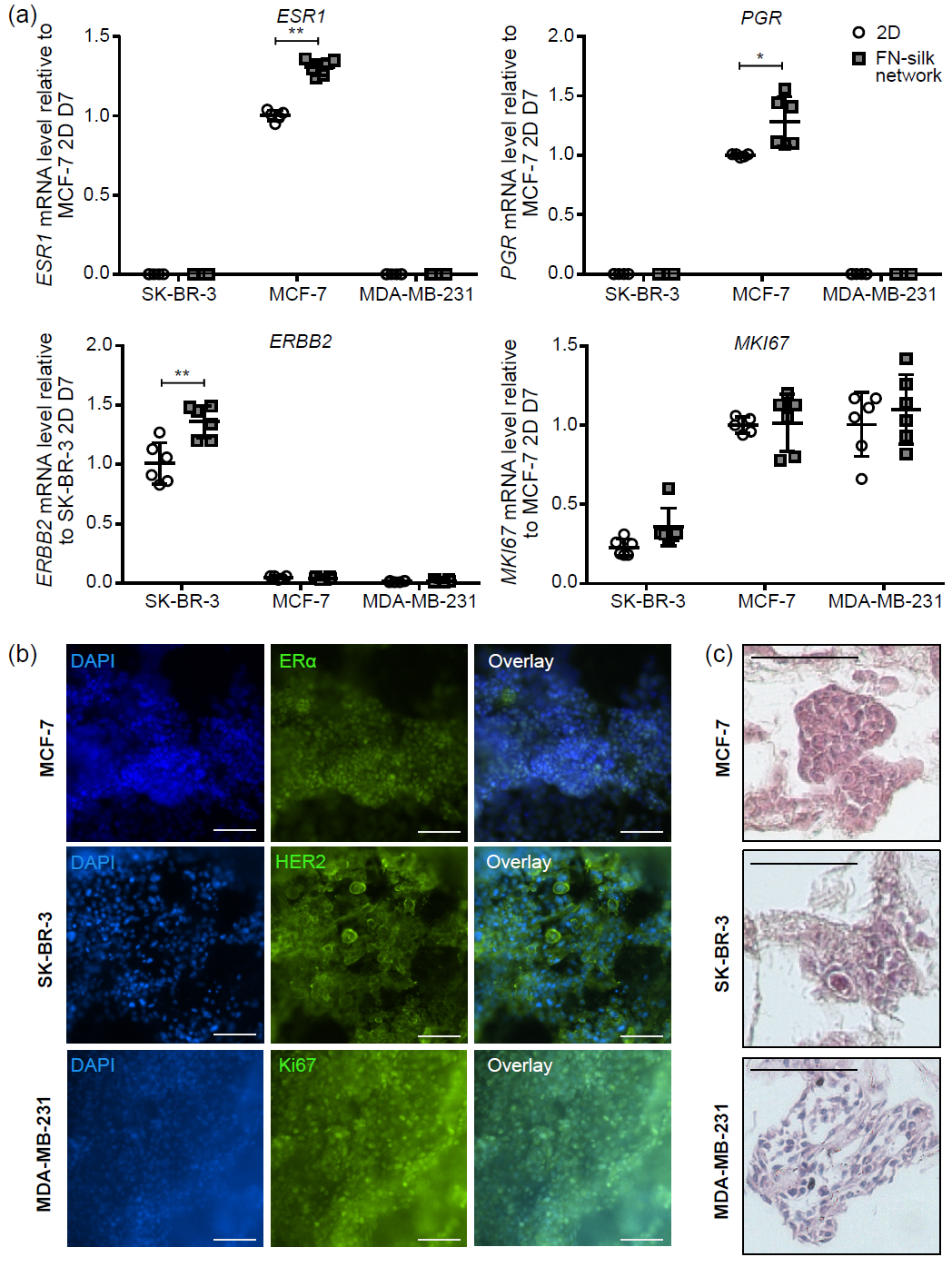
Breast cancer markers and morphological appearance of cell lines cultured in FN-silk networks. (a) Gene expression levels of the canonical breast cancer markers *ESR1, PGR, ERBB2*, and *MKI67* were measured on day 7 in SK-BR-3, MCF-7, and MDA-MB-231 cells kept in culture in 2D or FN-silk network. Values are represented as fold-change of the mean ± SD (n=6). Fold-change was calculated comparing to the control MCF-7 2D on day 7 for *ESR1, PGR*, and *MKI67* and to control SK-BR-3 2D on day 7 for *ERBB2. *P <* 0,05; *\*\*P <* 0,01. (b) Immunofluorescence staining was done on MCF-7, SK-BR-3, and MD A-MB-231 cells cultured in FN-silk network for 7 days. MCF-7 was stained for ERa (green), SK-BR-3 for HER2 (green), and MDA-MB-231 for Ki67. Cell nuclei were stained with DAPI (blue), and an overlay image was created. Scale bar: 100 μm. (c) Hematoxylin and eosin staining on sections of MCF-7, SK-BR-3, and MDA-MB-231 cells cultured for 7 days in FN-silk network. Scale bar: 100 μm.

To confirm the expression at the protein level of ER*α* and HER2, we performed staining on whole networks after seven days of culture. The MCF-7 cells showed a nuclear signal for ER*α*, while the SK-BR-3 stained positively for HER2 (Fig. 3b). As expected, the triple-negative MDA-MB-231 was negative for both markers (Supplemental Fig. 1b) while abundantly expressing Ki67 (Fig. 3b). Lastly, we performed hematoxylin and eosin (H&E) staining on sections of the networks (Fig. 3c), detecting a distinct morphology for each cell line. For instance, MDA-MB-231 spread without a clear spatial reorganization, SK-BR-3 tended to create loose cell clusters, and MCF-7 arranged themselves into clusters with a diameter of approximately 100 μm (Fig. 3c).

### Culture in FN-silk networks modulates gene expression and confers signatures of invasiveness

Growth in a 3D microenvironment is known to affect the gene expression profile of cells and render them closer to actual tissues.^22,23^ Processes such as adhesion, migration, angiogenesis, and epithelial-mesenchymal transition (EMT) play a role in tumor development and invasiveness. To test if growth in FN-silk networks affects these biological mechanisms, we performed RT-qPCR to investigate the expression of 34 representative genes. The results revealed significant changes for 12 of the 34 targets (Supplementary Fig. 2a and 2b). Growth in FN-silk networks modulated several cell adhesion markers in MCF-7 (*i*.*e*., *CDH2, JAM2, CLDN3)* and promoted the expression of *CD44*, a glycoprotein involved in cell-cell interactions, cell adhesion, and migration, in both MCF-7 and SK-BR-3. Moreover, MDA-MB-231 cells grown in FN-silk networks had high cancer stem cell markers (*i*.*e*., *NANOG, OCT4*, and *SOX2)*, accompanied by an up-regulation of genes associated with basement membrane degradation and invasiveness (*i*.*e*., *HIF1α, ITGAV*, and *MMP14)*.

To obtain the complete global transcriptome changes driven by culture in FN-silk networks, we decided to use RNA-sequencing (RNA-seq). For this experiment, we focused on the two cell lines MCF-7 and MDA-MB-231. This choice was motivated by the observed robust metabolic rate and the high number of modulated transcripts associated with culture of these two lines in FN-silk. For the RNA-seq, we compared three conditions, being 1) cells directly lysed after trypsinization from the flask at day zero, referred to as the control flask, 2) cells cultured for seven days in FN-silk network, and 3) cells that were grown for seven days on a flat tissue-culture treated plate, as illustrated in the cartoon of Fig. 4a.

**Fig. 4.**
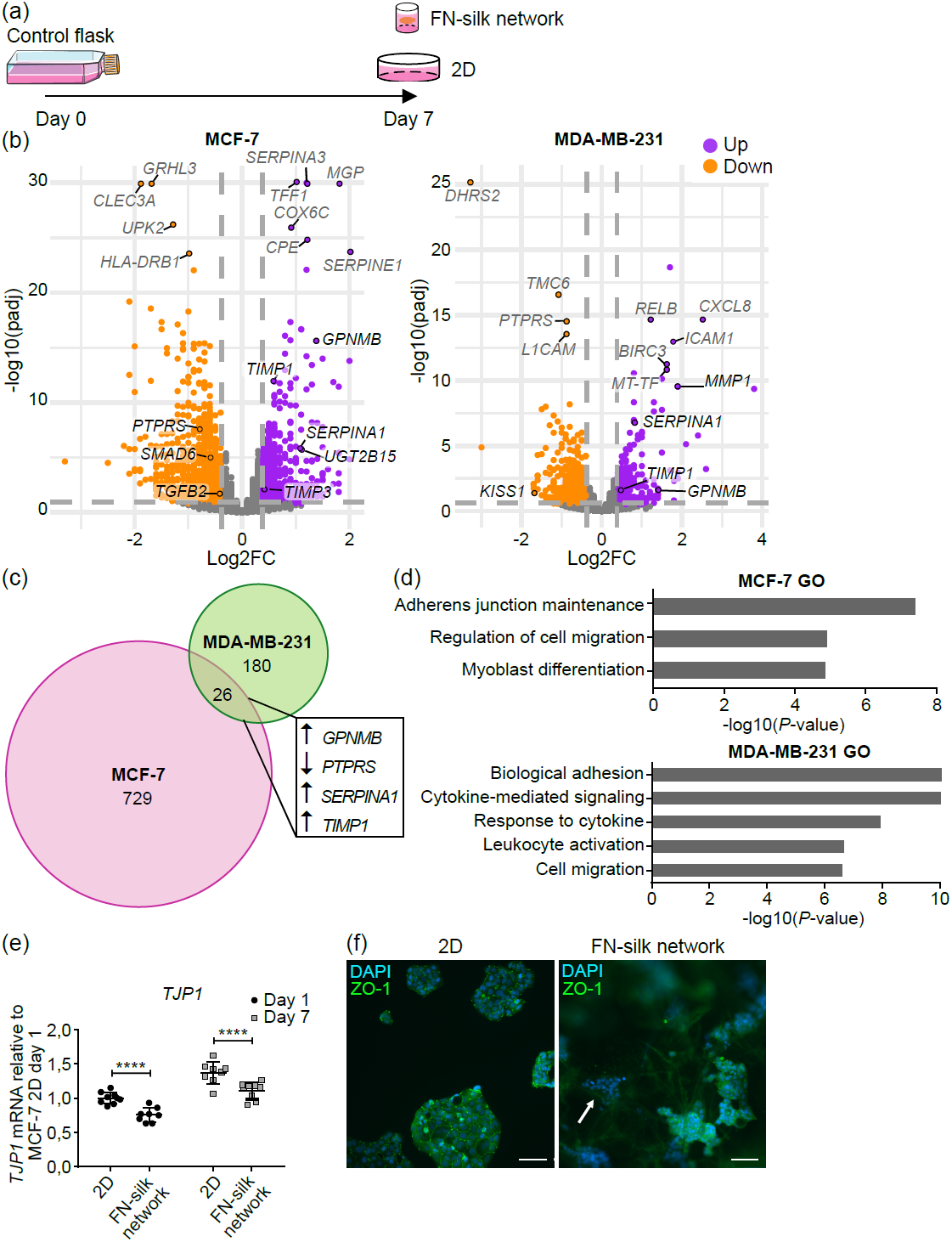
Impact of FN-silk network on global gene expression levels and ZO-1 target validation. **(a)** Schematic illustrating the three conditions compared for the transcriptomic experiment, being i) cells lysed from the control flask at day 0, ii) cells lysed after 7 days in culture in 2D, or iii) 7 days in culture in FN-silk network, (b) Volcano plots of MCF-7 and MDA-MB-231 transcriptomes after 7 days in culture in FN-silk network compared to their counterparts grown in 2D. The x-axis corresponds to the log2FC and the y-axis to the negative loglO of the adjusted P-value. Transcripts with an adjusted *P-* value < 0,05 are shown in purple if log2FC > 0,38 and in orange if log2FC < −0,38. The threshold values are represented with thicker dark grey lines. Transcripts not meeting the criteria are depicted in grey. The most significantly regulated genes are indicated in dark grey, whereas selected targets relevant to the pathophysiology of cancer are indicated in black, (c) Proportional Venn diagram representing the number of significantly differentially expressed genes due to culturing of MCF-7 (pink) and MDA-MB-23 1 (green) in FN-silk network. 26 transcripts are commonly regulated in both cell lines. Among these, genes with key roles in breast cancer are indicated (*i*.*e*., *GPNMB, PTPRS, SERPINA1, TIMPP)*. (d) Gene enrichment analysis of the FN-silk network signature in MDA-MB-231 and MCF-7. Web-Gestalt was used to explore the gene ontology terms associated with the FN-silk network signature. The bars represent the negative log 10 *(P-*value) of enriched terms, indicating the significance of the association between the gene list and an indicated ontology term, (e) mRNA levels of tight junction protein-1 *(TJP1)*, a significantly decreased gene in MCF-7 grown in FN-silk network compared to cells kept in 2D. Values are represented as fold-change of the mean ± SD (n=3 with three technical replicates). Foldchange was calculated compared to the control MCF-7 2D on day 1. For the statistical analysis, 2-way ANOVA was performed, *\*\*\*\*p <* 0,0001. (f) *TJP1*-encoded protein zonula occludens-1 (ZO-1, green) and nuclei (DAPI, blue) were stained in MCF-7 cultured in 2D or FN-silk network for 7 days. The arrow points to a cell cluster in which ZO-1 was not detected. A representative overlay image is shown. Scale bar: 100 μm.

A principal component analysis (PCA) was conducted to detect eventual outliers and to evaluate the effect of cell type (*i*.*e*. MCF-7 and MDA-MB-231), culturing condition (*i*.*e*. 2D or 3D), and time (*i*.*e*. day 0 and 7) on the results. The first exploratory PCA analysis took into account the results from both cell lines and revealed the cell type as the factor accounting for 99% of the variance (Supplemental Fig. S3a). This data confirmed a major difference between the two breast cancer subtypes. Considering such observation, we processed the datasets generated using the two cell lines independently. These PCAs showed that for both MDA-MB-231 and MCF-7, the cultivation of cells in FN-silk network caused the greatest variance, while time had only a minor effect (Supplemental Fig. S3b, c). Additionally, we observed a greater effect caused by FN-silk for MCF-7 (62%) than for MDA-MB-231 (46%).

Culture in FN-silk network led to 755 differentially expressed genes (DEGs) in MCF-7 and 206 in MDA-MB-231, which had a similar distribution between up- and down-regulated, as shown by the volcano plots (Fig. 4b, full list of DEGs is available in Supplemental Table 1). While 26 DEGs were shared between the two cell lines (Fig. 4c), only 17 responded in a similar fashion. Among these 17 genes, there were genes involved in cancer progression, for instance, the down-regulated tumor suppressor *PTPRS*^24^ and the up-regulated *SERPINA1* whose overexpression was reported to increase invasiveness,^25^ *GPNMB*, which induces stem-cell-like properties in cancer cells,^26^ and *TIMP1* a promoter of tumorigenesis.^27^ These 17 genes were also tested in SK-BR-3 cells, showing a conserved decrease in *AP1G2 and PTPRS*, as well as an increase in *TIMP1* (Supplementary Fig. 4b).

**Table 1.**
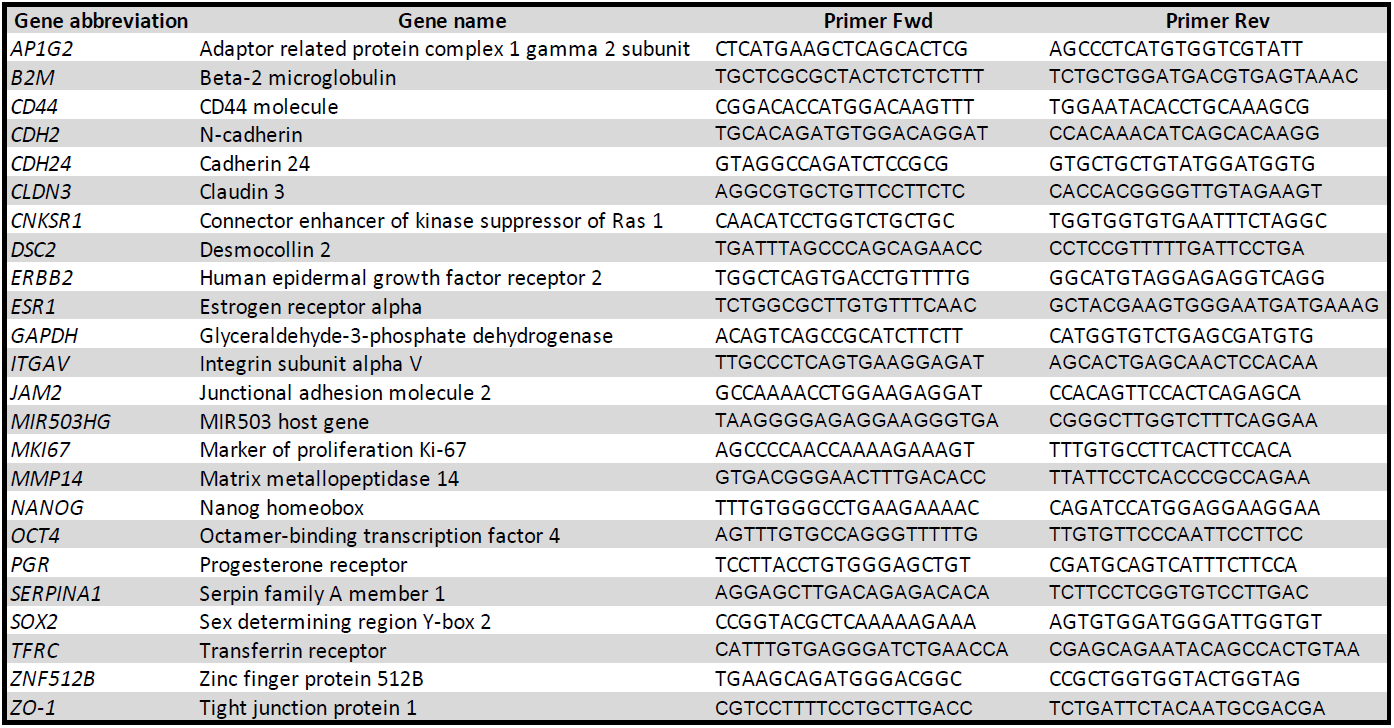
RT-qPCR primers. List of the human primers used in this study completed of gene abbreviation, gene name, and forward/reverse sequences.

Gene ontology analyses revealed cell-specific signatures, such as inflammation in MDA-MB-231 and myoblast differentiation in MCF-7, as well as common biological processes of migration and adhesion. Additionally, to investigate whether the genes modulated by culture in FN-silk networks had a profile relevant to breast cancer biology, we performed a gene set enrichment analysis (GSEA) against a functional database built on the genes of the Nanostring nCounter® Breast Cancer 360 panel, which includes 758 genes that cover established breast cancer diagnostic and research signatures (e.g. PAM50) as well as key pathways of the tumor and its microenvironment. This data exploration showed a significant enrichment score for the genes in MCF-7, highlighting that cultivation in FN-silk networks affects the expression of targets important for breast cancer biology (Supplementary Fig. 4a).

To investigate which transcription factors (TFs) were mediating the observed changes, we performed two bioinformatic predictions. The first was done using i-*cis*Target, an integrative method that identifies shared regulatory regions in the promoters of sets of DEGs.^20^ The results revealed RELA/P65, a subunit of the NFKβ inflammatory complex, as the main responsible for the changes observed in MDA-MB-231, corroborating the gene ontology results and suggesting a higher inflammatory response in MDA-MB-231 cultured in FN-silk networks than in 2D. In MCF-7, the promoters of the DEGs were mostly enriched for two TFs, being 1) transcription factor AP-2 alpha (TFAP2A), which is known to play a role in EMT,^28^ and 2) the TEA domain family members (TEADs), who support cancer progression by promoting genes related to proliferation.^29^

For the second bioinformatic prediction, we took advantage of a complementary approach based on prior knowledge of expected effects between transcriptional regulators, not exclusively TFs, and their target genes stored in the Ingenuity® Knowledge Base.^21^ While the results confirmed NFKβ as the most activated regulator responsible for the changes of MDA-MB-231, they highlighted nuclear protein 1 (NUPR1), previously associated with breast cancer metastasis, as the most significantly activated regulator in MCF-7 (full list available in Table 2). In addition to being predicted as activated, *NUPR1* also had an increased mRNA level in MCF-7 grown in FN-silk. NUPR1 has emerged as a repressor of ferroptosis, a type of iron-dependent regulated cell death.^30^ Since a recent study suggested a new tumor-mediated control of iron permitted by the 3D tumor architecture,^31^ we explored the DEGs in MCF-7 for targets related to iron storage. We found that culture in FN-silk increased both light *(FTL)* and heavy *(FTH1)* chains of Ferritin, the major intracellular iron storage protein.

**Table 2.** Transcriptional regulators. List of the upstream regulators predicted to mediate the gene expression changes observed in MCF-7 and MDA-MB-231 following culture in FN-silk networks.

| MCF-7 |  |  |  |
| --- | --- | --- | --- |
| Transcriptional regulator | Predicted state | Activation z-score | *P*-value |
| NUPR1 | Activated | 3,3 | 0,0175 |
| ESR1 | Activated | 3,1 | 4,1E-0,5 |
| WBP2 | Activated | 2,4 | 0,0204 |
| TCF4 | Activated | 2,3 | 5,23E-0,4 |
| RAD21 | Inhibited | -2,0 | 5,87E-0,4 |
| NANOG | Inhibited | -2,1 | 7,35E-0,3 |

| MDA-MB-231 |  |  |  |
| --- | --- | --- | --- |
| Transcriptional regulator | Predicted state | Activation z-score | *P*-value |
| RELA/NFKB | Activated | 2,9 | 4,5E-11 |
| SREBF1 | Activated | 2,9 | 1,5E-05 |
| NFAT5 | Activated | 2,8 | 1,0E-05 |
| EGR1 | Activated | 2,8 | 1,72E-0,5 |
| NCOA2 | Activated | 2,4 | 8,51E-0,4 |
| ECSIT | Activated | 2,4 | 1,10E-0,7 |
| ATF4 | Activated | 2,4 | 7,39E-0,4 |
| STAT1 | Activated | 2,4 | 1,25E-0,2 |
| BHLHE40 | Activated | 2,4 | 5,8E-04 |
| HMGB1 | Activated | 2,4 | 1,73E-0,7 |
| FOXL2 | Activated | 2,2 | 2,71E-0,4 |
| EZH2 | Activated | 2,2 | 6,13E-0,5 |
| GFI1 | Inhibited | -2,0 | 1,7E-04 |
| BCL3 | Inhibited | -2,3 | 6,3E-06 |
| SATB1 | Inhibited | -2,4 | 3,0E-03 |
| ZFP36 | Inhibited | -2,4 | 1,7E-06 |

Tight junctions have a vital role in maintaining cell-to-cell integrity, and the loss of cell cohesion can lead to invasion and, thus, metastasis of cancer cells.^32^ Tight junction protein 1 *(TJP1)* is considered a tumor suppressor, and its downregulation has been associated with invasive features of cancer. We identified *TJP1* among the adhesion genes with decreased expression in MCF-7 grown in FN-silk networks. We confirmed that the decrease of *TJP1* can be detected already after 24 hours and is maintained over time (Fig. 4e). Afterward, we performed staining for the protein coded by *TJP1*, zonula occludens-1 (ZO-1), revealing that MCF-7 cultured in 2D formed monolayer clusters in which ZO-1 was always detectable, whereas in FN-silk networks some clusters did not express ZO-1 (Fig. 4f and Supplementary Fig. 4c).

### FN-silk networks as a flexible system to grow physiologically relevant cancer cells

In the last part of the study, we investigated if it is possible to use the FN-silk networks for the culture of novel breast cancer cell lines as well as primary cells obtained from fresh human breast tumors.

First, we tested if FN-silk networks could support the growth of the Wood and PB cells, two lines with phenotypic properties which faithfully reproduce the original cancer tissues.^33^ These cells complement canonical immortalized cells by better representing the variability observed among cancer patients. The comparison of breast cancer markers between Wood, PB, and the canonical SK-BR-3, MCF-7, and MDA-MB-231, showed how the ER*α*+ Wood cells have much lower levels of ER*α* than the classically used MCF-7, whereas PB resembles a triple-negative subtype, but with lower proliferation rate than the MDA-MB-231 (Supplementary Fig. 5).

Staining for the cytoskeleton protein F-Actin demonstrated that Wood and PB cells could homogeneously spread in FN-silk networks (Fig. 5a). The results also indicated a cell-specific morphology. For instance, Wood cells tended to reorganize themselves into clusters, and PB cells created a more continuous layer of cells. The distinctive cellular architecture was confirmed by H&E staining. Additionally, we tested if any of the 17 commonly regulated genes in MCF-7 and MDA-MB-231 were modulated in Wood and PB. The results showed that for Wood cells, culture in FN-silk networks caused a decrease in *AP1G2, CNKSR1*, and *ZNF152B*, as well as increased *SERPINA1*. For PB cells, we detected a significant down-regulation of *AP1G2, CDH24*, and *MIR503HG* (Fig. 5b).

**Fig. 5.**
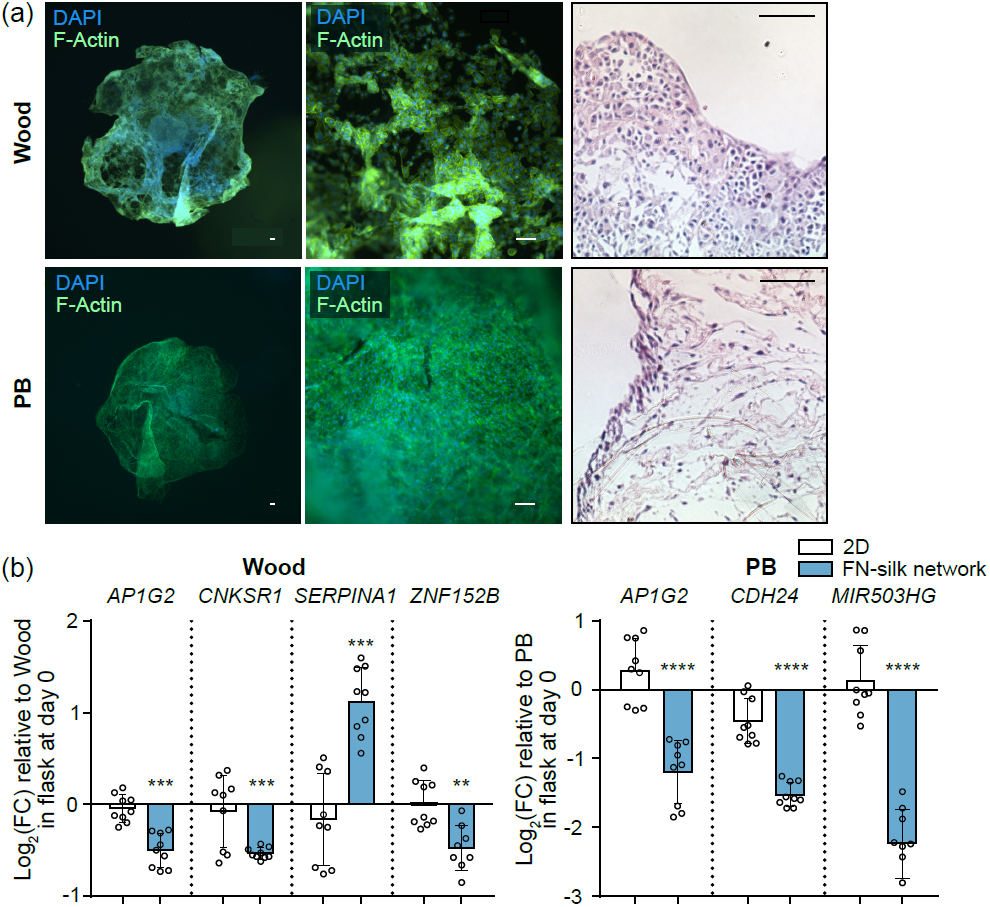
Wood and PB breast cancer cells grow in FN-silk networks. (a) Wood and PB breast cancer cells cultured for seven days in the FN-silk network were stained for actin filaments (F-Actin, green) and nuclei (DAPI, blue). Overlay images of pictures taken with 2X and 10X magnification are shown for both cell lines. Scale bar: 100 μm. The figures on the right side of the panel show hematoxylin and eosin staining on sections of Wood and PB cells cultured for 7 days in FN-silk network, 10X magnification. Scale bar: 100 μm. (b) Transcript levels of genes commonly regulated by growth on the FN-silk network were measured on Wood and PB harvested from the control flask on day 0 or from cells grown in 2D or FN-silk network for 7 days. Values are represented as log2 fold-change of the mean ± SD (n=9). Fold-change was calculated compared to the control flask on day 0. For the statistical analysis, 2-way ANOVA was done. *\*P <* 0,05; *\*\*P* < 0,01; *\*\*\*P <* 0,001; *\*\*\*\*p<* 0,0001.

Finally, we examined the potential of FN-silk networks to grow primary cells obtained via superficial scraping of tumor material following surgical removal.^34^ For each sample, the tissue was subjected to enzymatic dissociation. The cell suspension was then seeded on tissue-culture-treated 96-wells or mixed with FN-silk. To compare the viability of the patient-derived cells in 2D and FN-silk networks, live/dead staining was performed after seven days of culture. The data revealed alive cells across the FN-silk network (Fig. 6a-I). Areas with a high density of cells were also detected (Fig. 6a-II and -III), suggesting that even incompletely dissociated tissue was kept alive in the network. Such observation was in contrast with what was seen in the 2D control, where floating cell clusters were visible directly after seeding (Fig. 6b) but lost during the routinely performed medium change.

**Fig. 6.**
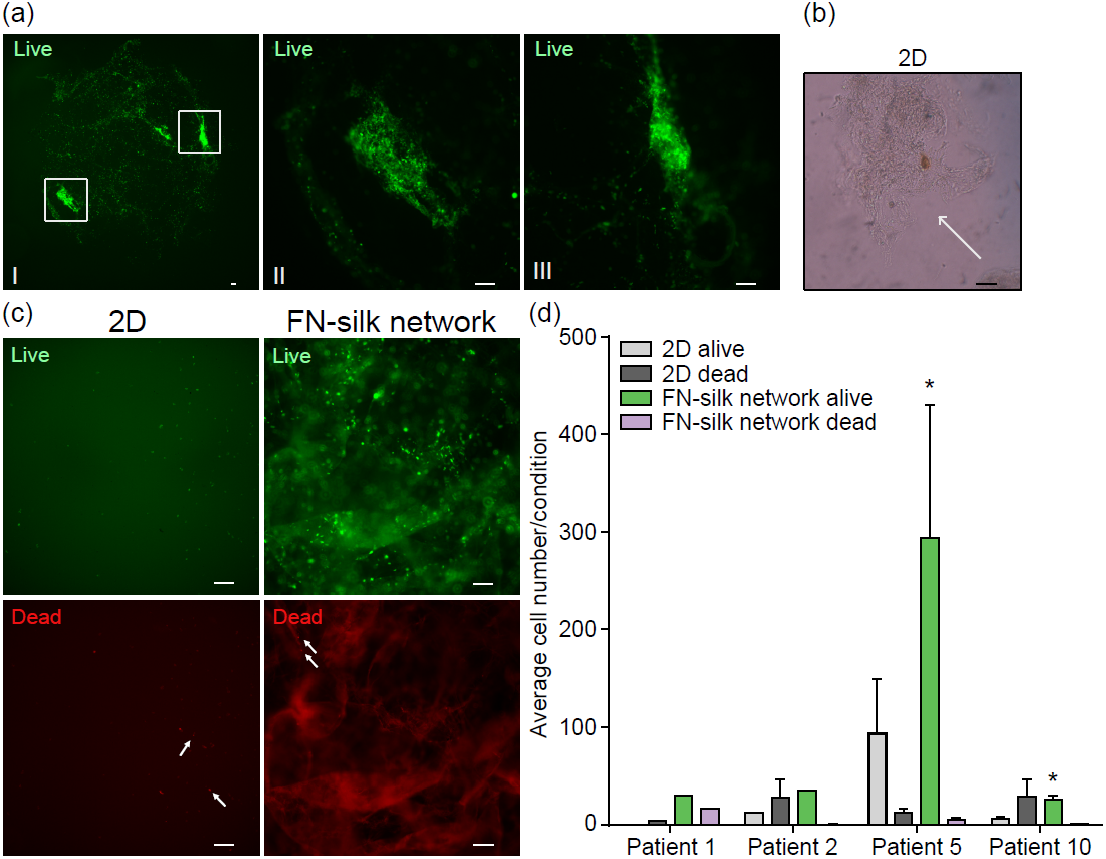
FN-silk network promotes viability of cells dissociated from patients’ tumors. Live/dead assay is done on FN-silk networks containing cells obtained from breast cancer tissue and cultured for seven days, (a) Live channel image of a FN-silk network scaffold generated using cells from patient 5. Picture (I) was taken with a 2X magnification, while (II) and (III), which show areas with abundant cell clusters, were obtained with a 10X magnification. Scale bar: 100 μm. (b) Brightfield image of a cell cluster floating in a 2D well, following the dissociation of the original tumor material. 10X magnification, scale bar: 79 μm. (c) Representative pictures of live (green) and dead (red) staining of cells grown in 2D and FN-silk network for 7 days, 10X magnification. The white arrows point to examples of dead cells. Scale bar: 100 μm. (d) Quantification of alive and dead cells from four biological replicates. No pink bar is visible for the biopsies corresponding to patients 2 and 10 because no dead cells were observed in the FN-silk network condition. For the statistical analysis, 1-way ANOVA was done. *\*P <* 0,05.

After the dissociation step, we generally found a low number of cells, with the enzymatic dissociation leading to approximately 20.000 cells per sample. Because of the low cell number constraint, we were able to perform quantifications of live and dead cells for only four of the ten biopsies. This was done in ImageJ by counting the number of cells in three different regions of each sample from the live (green) and dead (red) channels (Fig. 6c). Notably, FN-silk has an autofluorescence emission in the red channel, which causes a red background signal, examples of dead cells in FN-silk network are indicated with white arrows (Fig. 6c). The results of this quantification revealed a higher number of cells in FN-silk than in 2D, with more dead cells visible in 2D than in FN-silk networks (Fig. 6d). Overall, these results indicate that the FN-silk scaffold can support patient-derived cell growth more efficiently than conventional 2D culture.

## 4. Discussion

Our findings demonstrate the possibility of growing immortalized cells as well as primary cells obtained from the superficial scraping of fresh tumors in floating networks made of FN-silk. Such a model shows an advantage over gel-based and scaffold-free alternatives since it recapitulates not only cell-cell but also cell-extracellular matrix (ECM) interactions. Additionally, while existing scaffold-based options often suffer from batch-to-batch variability^12^ and low seeding efficiency,^35^ the herein-described FN-silk has a completely controlled composition and ensures optimal seeding distribution thanks to its self-assembly properties. These strengths are complemented by FN-silk being a completely defined protein, cell compatible,^15,17^ and easy to handle at room temperature for the time required by the described protocol.

Previously, we showed that pipetting air bubbles into a solution of FN-silk leads to the self-assembly of FN-silk sheets around each air bubble, which, when bursting, yields a network where cells can stretch out and proliferate.^16^ These silk networks had been prepared anchored to the bottom of a tissue-culture plate. We here adapted the published protocol introducing three main novelties to the method, being; a) the use of immortalized and primary human breast cancer cells, b) a step for fast removal of air bubbles from the scaffold to better mimic physiological conditions, and c) the culturing of a floating network in which all the sides are equally exposed to the medium.

Several publications have reported that cells proliferate slower in 3D than 2D.^36^ In our study, metabolic activity measurements, commonly used to infer the proliferation rate of cells, only partly supported the existing literature. For instance, while the results obtained with SK-BR-3 and MDA-MB-231 confirmed previous data, MCF-7 had higher metabolic activity in FN-silk than 2D. The precise mechanism leading to this difference is unclear. However, we speculate it may be related to the cell-specific modality of reorganizing into space. As we estimated the area of each network to be about 40% smaller than the one of a 96-well, such a factor might have limited the proliferation of cells spreading without complex cell-cell morphologies (*i*.*e*. SK-BR-3 and MDA-MB-231) and, on the contrary, favored MCF-7, which created multiple compact cell clusters into the FN-silk construct.

Breast cancer patients are routinely stratified based on their distinct expression of markers. Since prognoses and treatments are tailored for each subtype, representative cell lines must recapitulate the features of the various types. We showed that culture in the FN-silk network does not lead to a loss in breast cancer markers. Additionally, we proved how each cell line developed a distinctive morphology when cultured in FN-silk, consistent with what was observed in Matrigel-based models.^37^

Studies using spheroids and gel-based scaffolds have demonstrated that growth in environments more complex than classical flat surfaces influences gene and protein expression.^22^ Herein, we investigated the transcriptional changes occurring when cancer cells are maintained in silk-based networks. Our sequencing data revealed cell-type-specific gene regulation driven by these culture conditions. While the affected genes largely differed among the two cell lines, the analysis revealed that many were related to cell adhesion and migration. Both MCF-7 and MDA-MB-231 had upregulated expression of genes promoting invasiveness (*SERPINA1*^25^), extracellular matrix remodeling (*TIMP1*^27^), and tumorigenicity (*GPNMB*^26^). N-cadherin, a protein mediating cell-cell contact, often upregulated in aggressive cancers, and reported to promote migratory behaviors in a collagen-based 3D model,^38^ was highly expressed in cells cultured in FN-silk. Growth in the FN-silk network led to a decrease in ZO-1, a protein that anchors tight junctions to the actin filaments of the cytoskeleton and whose down-regulation correlates with invasiveness.^39^ Overall, these results suggest that our scaffolds efficiently mimic the ECM, offering an environment where tumor cells can attach, modify, and migrate through the remodeled matrix.

Some of the changes observed in the triple-negative cells comprised a) an increase in cancer stem cell markers, similar to what was found in collagen and spheroid models,^40^ b) higher expression of the hypoxia-inducible factor *HIF1α*, which promotes angiogenesis and capability to metastasize^41^ and c) an NFκB-mediated inflammatory signature. NFκB, the main driver of cytokine expression in immune cells, was reported to promote the secretion of pro-inflammatory cytokines and contribute to aggressive tumoral features in triple-negative cells.^42,43^ Our data suggest that culture in FN-silk could trigger a high level of pro-inflammatory signaling. Since NFκB inhibitors are emerging as drugs to treat triplenegative cancers, our model could be used to study such medications.

Most of the literature comparing gene expression changes in 2D as opposed to 3D models has focused on a small set of genes. To our knowledge, only two studies considered the global transcriptional changes of MCF-7 grown in 3D, thus being somewhat comparable to our investigation. Blanchette-Farra’s research analyzed transcriptomic variations in MCF-7 cultured in three spheroid models.^31^ They generated spheroids using; a) plates coated with Poly(2-hydroxyethyl methacrylate), b) ultra-low attachment plates, and c) methylcellulose as an aggregating agent. Spheroids from these conditions had similar transcriptional changes (represented by 770 differentially expressed genes (DEGs)). In another study, Wulftange *et al*. maintained MCF-7 in Matrigel mixed with fibronectin solution and identified 3156 DEGs.^44^ The signatures from the two published datasets shared 240 common DEGs. A similar low overlap is also present when we consider the FN-silk-dependent genes in MCF-7, leading to 74 genes in common with the spheroids^31^ and 208 with the Matrigel model.^44^ Our study and the previous two shared only 24 genes. Among these are a) an increase of *UGT2B15*, which detoxifies drugs and has been shown to influence carcinogenesis in hormone-dependent cancers,^45^ and b) a higher level of *FTL*, the light subunit of ferritin, a major intracellular iron storage protein. This comparison highlights the limited reproducibility among independent studies, which could be partially reduced using components with a highly defined composition and low batch-to-batch variability, such as FN-silk. Nevertheless, it also suggests the presence of conserved mechanisms triggered by three-dimensional settings, such as higher detoxification ability and, thus, drug resistance as well as modulation of iron-storage proteins.

Additionally, the study from Blanchette-Farra^31^ suggested a new tumor architecture-dependent increased ability of iron storage in 3D cultures, which reduces ferroptosis, an iron-regulated cell death. Consistently with their study, we found both light and heavy subunits of ferritin up-regulated in MCF-7 grown in FN-silk. Moreover, we observed an increased mRNA level of *NUPR1*, a transcription factor that can repress ferroptosis,^30^ and bioinformatically predicted its activation in MCF-7 grown in FN-silk. It is tempting to speculate that our model favors an increase in intracellular iron storage via ferritin that, together with the activated ferroptosis repressor NUPR1, may decrease ferroptosis and render MCF-7 cells more resistant.

While we could demonstrate that gene expression changes relevant to tumor development and aggressiveness are modulated in FN-silk networks, the precise mechanisms involved are still elusive. Future investigations should be performed to establish the contribution of the three-dimensional nature and of the RGD motif from fibronectin to the observed transcriptional changes.

Recapitulating the tumor microenvironment by mimicking the ECM and including cells representing tumor heterogeneity is key for creating *in vitro* models. FN-silk networks offer an ECM-like structure that could support the novel Wood and PB cells, which phenocopy the original primary tumors thanks to a defined and optimized medium.^33^ The protocol for FN-silk network formation was proved to be flexible and capable of sustaining cell viability even in the case of minimal initial material with slowly proliferating cells, such as the one obtained by superficial scraping of breast cancers.^34^ This is valuable since low cell number and slow proliferation are a challenge for spheroids formation.^46^ The detection of live cell clusters also confirmed the high adaptability of our 3D model. For instance, the enzymatic dissociation step had to be performed quickly to avoid harming the patient-derived cells. This technical limitation resulted in a suboptimal dissociation, as indicated by the presence of cell clusters lost during media exchange in 2D plates but maintained in FN-silk networks. The FN-silk scaffold’s exquisite ability to favor cell attachment led to generally higher viability of cells obtained via superficial scraping of breast tumor, offering encouraging proof-of-concept results. Future studies should investigate whether the ability of FN-silk networks to support cells independently of their proliferation ability and morphology could allow to co-culture different cell types and recreate the original tumor niche.

## Conclusion

In this study, we described a new method to generate floating networks of breast cancer based on FN-silk. We demonstrated that our model allows to culture well-established and novel cell lines. We performed the first investigation of gene expression changes driven by culture in FN-silk networks for the two highly used MCF-7 and MDA-MB-231, providing a solid foundation for future studies aiming to adopt FN-silk networks. The data highlighted how culture in FN-silk networks modulates adhesion, migration, and the expression of genes that play a role in tumor development. Overall, suggesting that our model recapitulates the tumoral features more faithfully than 2D. We also proved that patient-derived cells can be cultured in FN-silk networks, holding promise for the recreation of miniaturized tumoral niches.

## Contribution statement

**Caterina Collodet:** Conceptualization (equal); data curation (lead); formal analysis (lead); funding acquisition (equal); investigation (lead); methodology (equal); supervision (equal); validation (lead); visualization (lead); writing – original draft (lead). **Kelly Blust:** Data curation (supporting); formal analysis (equal); visualization (equal). **Savvini Gkouma:** Investigation (equal). **Emmy Ståhl:** Investigation (equal), formal analysis (equal), visualization (equal). **Xinsong Chen:** Resources (equal); methodology (supporting). **Johan Hartman:** Resources (equal); methodology (supporting). **My Hedhammar:** Conceptualization (lead), formal analysis (equal); funding acquisition (lead); methodology (equal); project administration (lead); resources (equal); supervision (equal); visualization (equal); writing – original draft (equal); writing – review and editing (lead).

## Acknowledgments

We thank the Bioinformatics and Expression Analysis (BEA) core facility at the Karolinska Institute in Huddinge for performing the RNA-seq experiment and initial data analysis. We thank Professor Johan Hartman and Dr. Xinsong Chen for kindly donating the SK-BR-3, MCF-7, and MDA-MB-231 and for providing the patient biopsies. We thank Cellaria for generously providing us with the Wood and PB cells as well as the Renaissance Essential Tumor Medium. We are grateful to Lars Hiertas Minne for supporting this work and Spiber Technologies AB for providing recombinant spider silk proteins.

## Conflict of interest declaration

M.H. has shares in Spiber Technologies AB, a company that aims to commercialize recombinant spider silk.

**Supplementary fig. 1.**
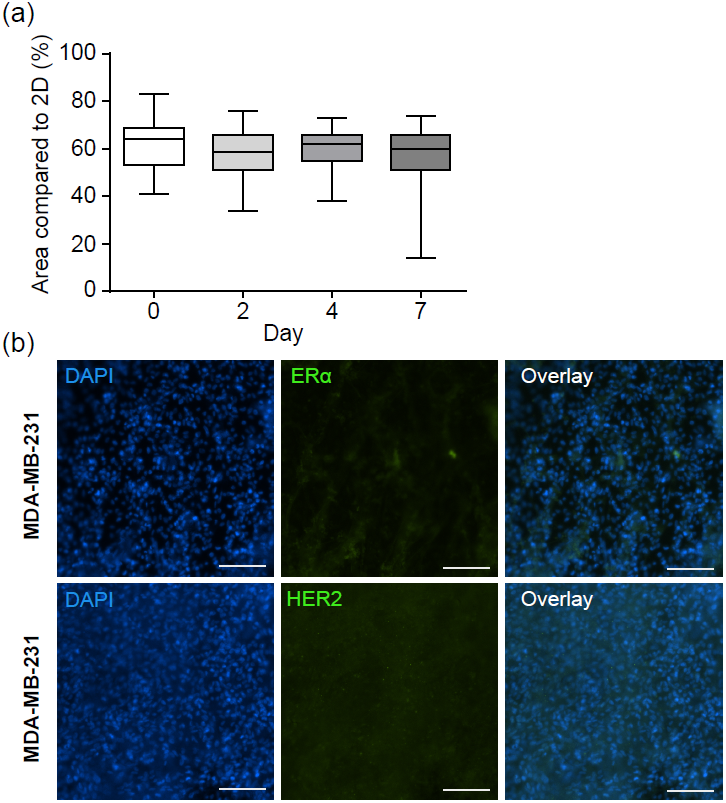
Surface area measurement time course of FN-silk network constructs. Box and whisker plots display the size of the FN-silk network areas as a percentage compared to the surface of a 96-well plate well (n=3 with 8 technical replicates per independent experiment), (b) Immunofluorescence staining for ER*α*, (green), HER2 (green), and nuclei (DAPI, blue) in MDA-MB-231 cells cultured in FN-silk network for 7 days. Single channels and overlay images are shown. Scale bar: 100 μm.

**Supplementary fig. 2.**
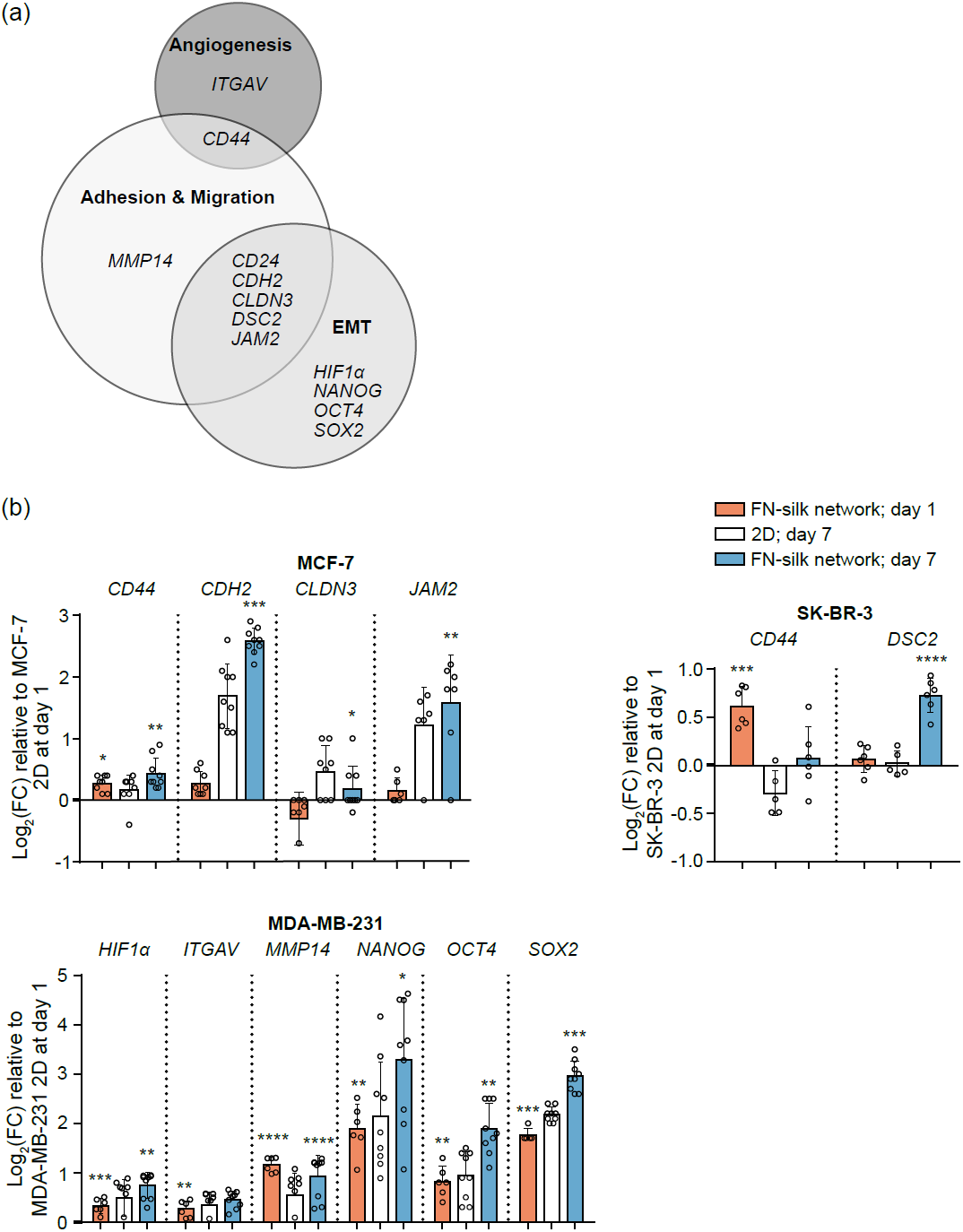
First RT-qPCR analysis of gene expression changes driven by FN-silk network. (a) Venn diagram representing the three biological processes, namely angiogenesis, adhesion and migration, and epithelial to mesenchymal transition (EMT), for which significantly regulated genes were identified, (b) Expression levels of the genes differentially expressed in MCF-7, SK-BR-3, and MDA-MB-231 due to cultivation in FN-silk network for seven days. Values are represented as log2 fold-change of the mean ± SD (n=6). Fold-changes were calculated compared to the control 2D on day 1. For the statistical analysis, 2-way ANOVA with Sidak correction was done. *\*P <* 0,05; *\*\*P* < 0,01; *\*\*\*p<* 0,001; *\*\*\*\*p<* 0,0001.

**Supplementary fig. 3.**
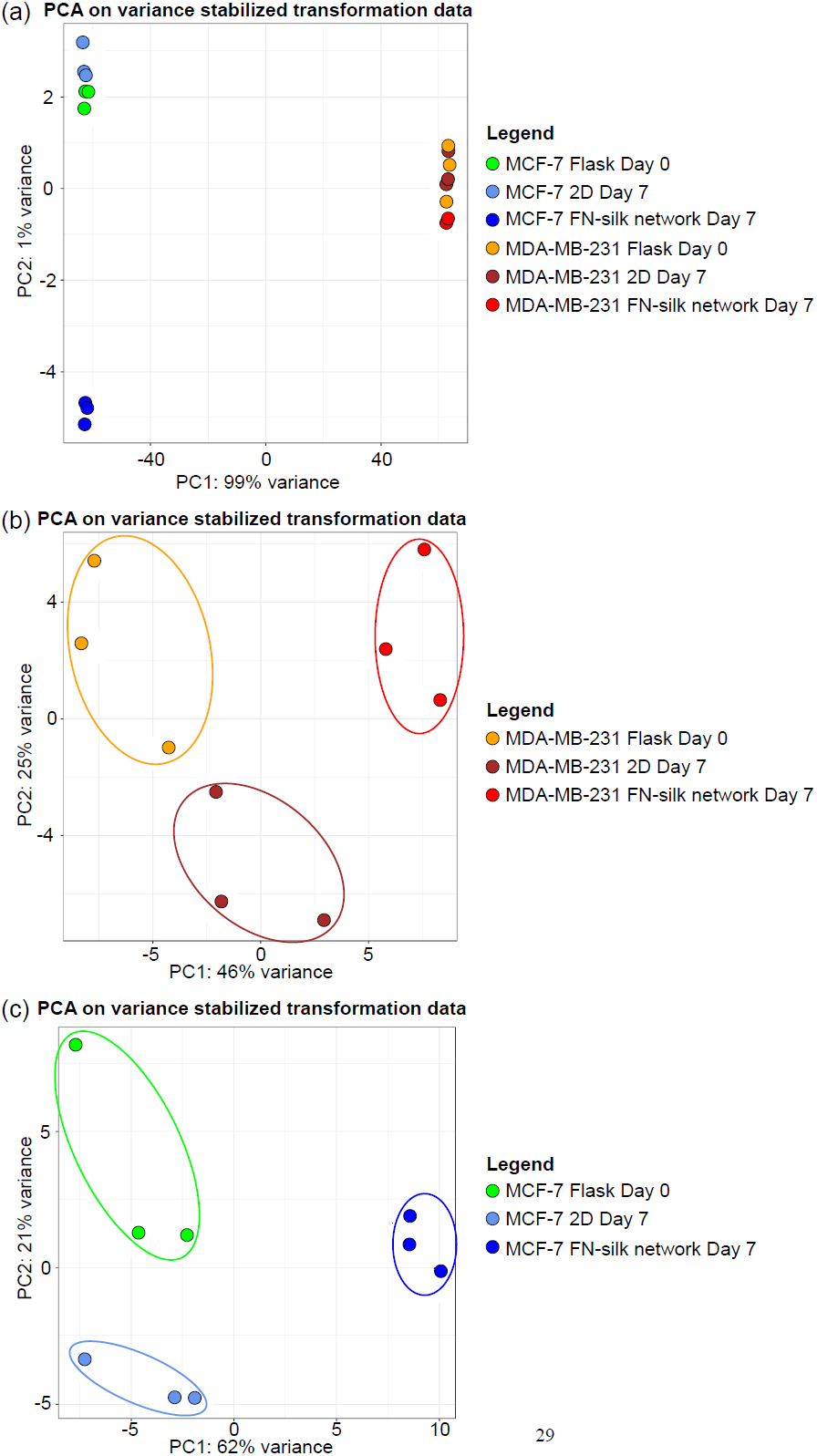
Principal component analysis (PCA) score plots. (a) PCA score plot based on the global gene expression data generated from comparing MCF-7 and MDA-MB-231 harvested at day 0 from the flask or at day 7 after culture in 2D or FN-silk network. The first two principal components (PC1 and PC2) explain 99% and 1 % of the total variance in the data and correspond to cell type and FN-silk network effect on MCF-7, respectively, (b) PCA score plot was obtained from the gene expression dataset solely of MDA-MB-231 samples. The FN-silk network after seven days in culture is responsible for 46% of the total variance in the data, as shown by PC1. (c) PCA score plot representing the MCF-7 subset of samples. The FN-silk network effect after seven days in culture accounts for 62% of the total variance in the data. Samples are depicted as indicated in the figure.

**Supplementary fig. 4.**
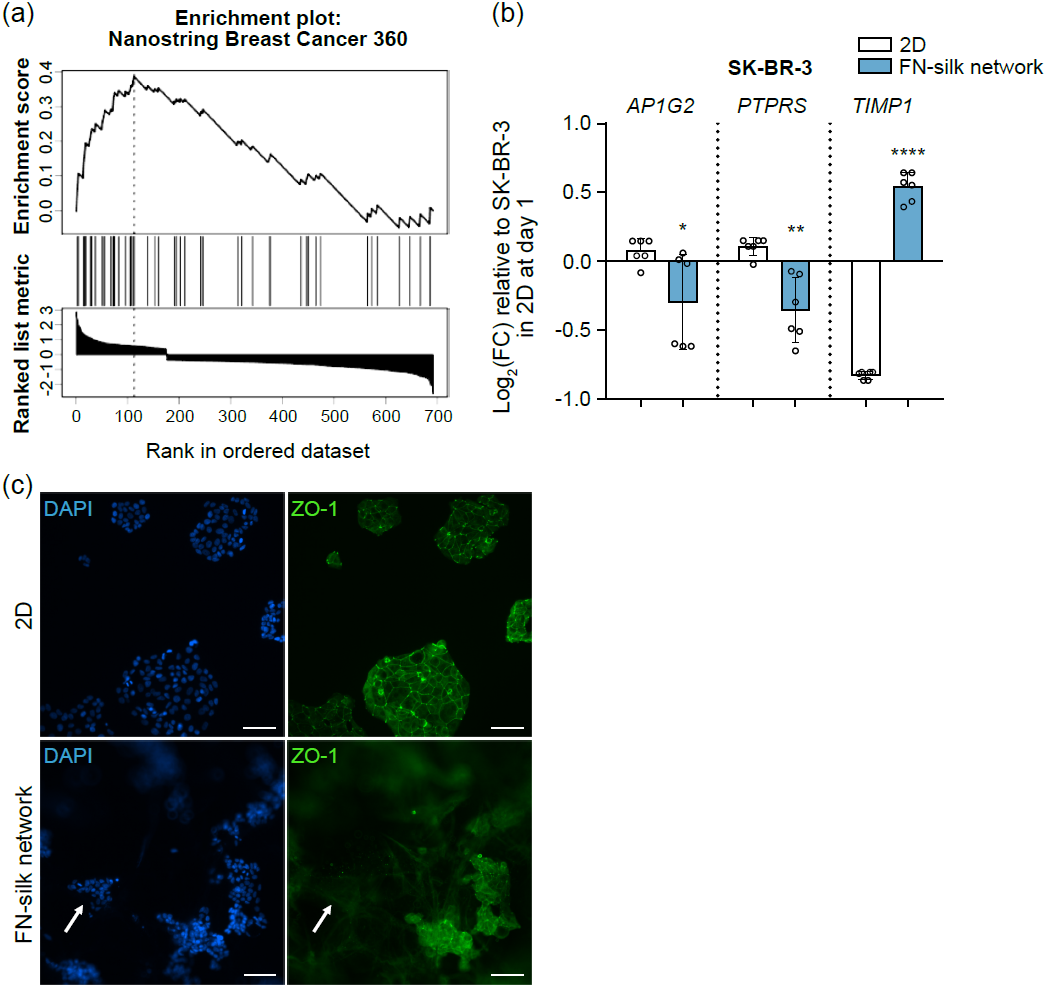
FN-silk network targets, further investigation at transcriptional and protein levels. (a) Gene set enrichment analysis (GSEA)-enrichment plot revealing a significant enrichment score when comparing the FN-silk network signature in MCF-7 with the gene panel of Nanostring Breast Cancer 360. (b) mRNA levels of genes commonly regulated by growth on FN-silk network were compared in SK-BR-3 grown in 2D or FN-silk network for 7 days vs at day 1 in 2D. Values are represented as log2 fold-change of the mean ± SD (n=2, three technical replicates). For the statistical analysis a t-test was done. *\*P <* 0,05; *\*\*P <* 0,01; *\*\*\*\*p<* 0,0001. (c) Staining of ZO-1 and nuclei done on MCF-7 cultured in 2D or FN-silk network for 7 days. Single channels are shown. Scale bar: 100 μm.

**Supplementary fig. 5.**
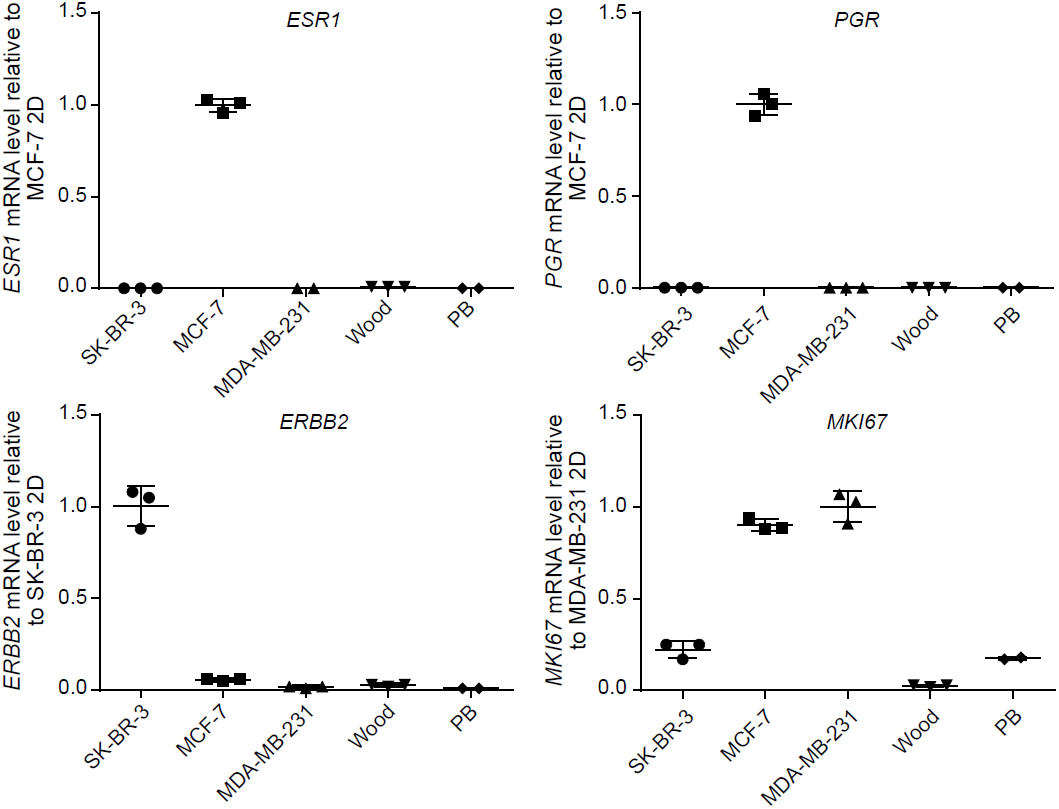
Breast cancer markers in Wood and PB cells. Gene expression levels of the canonical breast cancer markers *ESR1, PGR, ERBB2*, and *MKI67* were measured in SK-BR-3, MCF-7, MDA-MB-231, Wood, and PB cells kept in culture in 2D. Values are represented as fold-change of the mean ± SD (n=3). Fold-change was calculated comparing MCF-7 for *ESR1* and *PGR*, SK-BR-3 for *ERBB2*, and MDA-MB-231 for *MKI67*.

**Supplementary table 1**. List of differentially expressed genes in response to culture in FN-silk network.

## Supplementary materials and methods

### Cell culture

Three cell lines, MDA-MB-231, SKBR-3, and MCF-7, were obtained from American Type Culture Collection (ATCC) and cultured with a media mixture of DMEM low glucose (Thermo Fisher, 11574446), supplemented with 10% fetal bovine serum (Thermo Fisher, 16140-071) and 1% penicillinstreptomycin 10.000 U/ml (Thermo Fisher, 11548876). The Wood and PB cell lines, kindly donated by Cellaria, were cultured in RETM basal medium (Cellaria, CM-0001) supplemented with 5% heat-inactivated fetal bovine serum (Hyclone, SH3007103HI), 3% RETM supplement, 1% penicillinstreptomycin 10.000 U/ml, and Cholera toxin with a final concentration of 0,025 μg/ml (EMD Millipore, 227036). Regular passaging of the cell cultures with TrypLE (Thermo Fisher, 12605-028) was performed when cells reached a confluency of around 80%. The medium was changed every other day, and all cells were kept in a cell incubator at 37°C and 5% CO^2^. Cell counting was performed using a Bürker chamber (0,100 mm; 0,0025 mm^2^; 10^4^).

### Patient-derived cells isolation

In 2021, material from 10 breast cancer patients was collected at Karolinska University Hospital. All procedures were approved by the local ethical committee and performed under the ethical permit 2016/957-31 (Main application approval), 2017/742-32 (Amendment). Biopsies were obtained via superficial scrapings from breast tumors as previously described by Ma et al.^1^ and frozen in 10% DMSO solution. Cryovials were thawed at 37°C. Biopsy material was resuspended in warm DMEM/F-12, GlutaMAX medium (Thermo Fisher, 10565018) supplemented with 10% fetal bovine serum (FBS) and 1% penicillin-streptomycin 10.000 U/ml. The cell clumps were washed with a warm medium twice (370 g, 5 min). After the second wash, to enzymatically dissociate the cells, the cell pellet was incubated for 15 min in a cell incubator at 37°C and 5% CO^2^ in a medium containing prediluted dispase 1U/ml (Stemcell, 07923). During the incubation, cell clumps were gently mixed every 3 minutes. Following the incubation, a medium containing FBS was added to block the enzymatic digestion. The cell suspension was then filtered through a 100 μm strainer. Cells were counted and seeded on 96-well plate TC treated (Thermo Fisher, 174929) or in FN-silk networks as described in the next section. The number of seeded cells was adapted for each biopsy, depending on the initial number of available cells. One sample (patient 3) was excluded because all cells were found dead after the dissociation. For two samples (patients 2 and 4), only a couple of cells were detected in the Burker chamber during cell counting, with an approximated starting material between 13.000 to 20.000 cells.

### Alamar Blue proliferation assay

The metabolic rate of cells grown in tissue-culture treated 96-well plate or FN-silk network was measured using Alamar Blue cell viability reagent (Thermo Fisher, DALI 100) after 1, 3, and 7 days in culture. Briefly, after removal of the old culturing medium, 2D and 3D models were incubated for two hours at 37°C with 180 μl of Alamar Blue dye mixture diluted 1:10 in culturing medium. Three wells with only Alamar Blue were used as blank. After incubation, 100 μl of media were transferred to a new 96-well plate (Greiner, 655161), the plate was read using a plate reader (CLARIOstar, BMG Labtech) with excitation and emission wavelength of 544 nm and 595 nm, respectively. Fluorescence intensities of each well were obtained. For data analysis, the mean value of the blank was subtracted from each fluorescence intensity. For each condition, the average value and the standard deviation were calculated and plotted to compare 2D and FN-silk network cultures.

### Live/Dead viability assay

Cells obtained from the superficial scraping of tumors and cultured in 2D or FN-silk networks for 7 days were stained using the LIVEZDEAD™ viability/cytotoxicity kit (Invitrogen, L3224). After medium removal, 200 μl of a warm medium containing 0,05% calcein and 0,2% ethidium homodimer-1 (EthD-1) was added to each well or scaffold. Cells were kept in the incubator for 25 min, and afterward, pictures were taken with the ANDOR camera, Leica DMI6000B. Images were captured using the NIS elements BR software.

### Stereo microscope pictures of FN-silk network

A Nikon stereo microscope (SMZ 745T) was used to obtain macroscopic images of the FN-silk free-floating networks. The images were taken with a 1X magnification and analyzed with ImageJ to calculate the area of the FN-silk network model. The scale was set to 0,126 pixels/μm, based on the scale bar value obtained from the microscope.

### Brightfield microscopy of FN-silk network

An inverted phase contrast microscope (Nikon, TMS-F) equipped with a camera at 4X was used to obtain macroscopic pictures of the FN-silk network changes over the seven days of culture.

### Actin filaments staining of FN-silk network

Phalloidin staining was done to visualize actin filaments (F-actin) of cells grown in 2D and FN-silk network.

Samples were washed once with PBS and fixed with 4% paraformaldehyde (PFA) in PBS for 10 min at room temperature (RT) followed by three washes with PBS. The fixed cells were then permeabilized with 0,2% Triton X-100 in PBS for 15 min at RT. This was followed by two washes in wash buffer, Tween-20 (0,1 %) in PBS, and by blocking with 1 % bovine serum albumin (BSA) in PBS for 20 min at RT. Samples were then incubated for 40 min with phalloidin conjugated to Alexa Fluor 488 (Invitrogen, A12379) diluted 1:400 in PBS. An incubation step of 15 minutes with Sudan Black (Sigma Aldrich, 199664) 0,3% (w/v) working solutions dissolved in 70% ethanol was performed for FN-silk networks to quench the autofluorescence signal from the silk.^2^ Both models were washed three times with wash buffer before nuclear staining with 4’,6-Diamidine-2’-phenylindole dihydrochloride ((DAPI) Sigma Aldrich, 10236276001) diluted 1:1000 in PBS for 5 min. One final wash was done before imaging with the ANDOR camera, Leica DMI6000B. Images were captured using the NIS elements BR software.

### Immunofluorescence staining

Cells fixed as described in the previous section were permeabilized with 0,2% Triton X-100 in PBS for 15 min at RT, followed by blocking with 10% goat serum in PBS (*i*.*e*. block buffer) for 1 h. An overnight incubation in a humidified chamber at 4°C was done using primary antibodies diluted in a mixture of 50% block buffer and 50% wash buffer (0,1 % Tween-20 in PBS). The antibodies used were: HER2 (HP A, HPA001338, clone 181109,117P) diluted 1:250 and ER*α* (Invitrogen, MA5-14501, clone SP1) diluted 1:150. The following day, three 5 min washes with washing buffer were performed, followed by 1 h incubation at RT shielded from light with Alexa 488-labeled anti-mouse secondary antibody (Invitrogen, A32723) and Alexa 546-labeled anti-rabbit (Invitrogen, Al 1010) diluted 1:500 in a mixture of 50% block buffer and 50% wash buffer. After the incubation with secondary antibodies, FN-silk networks were washed once in wash buffer before an incubation step with Sudan Black (Sigma Aldrich, 199664) for 15 min. Both models were washed three times with wash buffer, and nuclei were stained with DAPI diluted 1:1000 in PBS for 5 min. Two final washes of 5 min in PBS were performed before imaging.

### Formalin-fixed paraffin-embedded (FFPE) samples preparation

FN-silk network samples were kept in culture for seven days. On day 7, samples were washed once with PBS and fixed in 4% PFA for 15 min at RT. After fixation, three washes of 5 min each in PBS were performed. To enable an easier localization of the samples during sectioning, FN-silk networks were incubated in a 1:1 solution of PBS: Mayer’s Hematoxilyn (Sigma, MHS32) for 30 s, followed by three 2 min washes with PBS. To maintain the integrity of the specimens, samples were pre-embedded in HistoGel (ThermoScientific, HG-4000-012). Briefly, a tube with solid HistoGel was heated to 60°C, and 150 μl of the liquid HistoGel was used to pre-coat a cryomold (10×10×5 mm). The FN-silk network was placed on top of the pre-coating and overlaid with an additional 200 μl of HistoGel. The cryomold was put on ice for 15 min for the Histogel to set. Pre-embedded samples were placed in a tissue embedding cassette and submerged in 70% ethanol. The samples were dehydrated using an automatized robot (Miles, Tissue-Tek V.I.P. E150/E300 Series). The program followed for dehydration consisted of two changes of 70% ethanol (30 min and 1 h), three changes of 95% ethanol (45 min, 45 min, and 1 h), and three changes of 99% ethanol (45 min and two of 1 h), two changes of xylene (1 h and 1 h 20 min), all performed at 40°C. These steps were followed by three changes of paraffin (two of 1 h and a third of 2 h) at 60°C. Finally, samples were embedded in paraffin using a tissue-tek embedding station (Microm, EC 350-1) and sectioned using a microtome (Microm, HM 360) to 12 μm thick sections.

### Hematoxylin and eosin staining

FFPE sections were deparaffinized with two 5 min changes of xylene (Sigma, 534056) followed by 5 min in 100% ethanol, 2 min in 95%, and 2 min in 70% ethanol. Sections were briefly washed in distilled water and stained for 2 min in Harris-modified hematoxylin solution (Merck, HHS32). Sections were then rinsed under tap water for 1 min, differentiated with two dips in 1% HO in 70% ethanol, rinsed with tap water, and blued in Scott’s tap water. Samples were then placed in 95% ethanol for 30 sec before counterstaining with 0,5% eosin for 3 min (Sigma, 861006), followed by rinsing in tap water. Dehydration was completed with sequential 1 min washes in 95% ethanol, 100% ethanol, and two changes of xylene. Sections were mounted with Permount mounting medium (VWR, 100496-552) and sealed with transparent nail polish.

### RNA extraction and cDNA synthesis

The medium was removed, and cells were washed once with PBS and lysed through the addition of cell lysis buffer for 2D (RNeasy minikit, QIAGEN, 74104) or FN-silk networks (RNeasy microkit, QIAGEN, 74034) to obtain a volume of 350 μl total. 2D cultures were collected by scraping the bottom of the 6-well plate with a cell scraper, the cells were collected in an Eppendorf tube. For the 3D cultures, five or three FN-silk networks were pulled together for lysis on day 1 and day 7, respectively. A syringe with a needle diameter of 0,4 μm was used to homogenize the samples via mixing 5-6 times, lysates were then transferred to an Eppendorf tube. Total RNA was extracted from lysed 2D models by using RNeasy minikit. For the FN-silk network, the silk debris was removed via centrifugation (12.000 g, 5 min), the supernatant was then processed using the RNeasy microkit. To measure the purity and quantity of RNA obtained, Nanodrop measurements were performed. Synthesis of cDNA was done using PrimeScript RT Kit (TaKaRa, RR037A) accordingly to the manufacturer’s instructions.

### RT-qPCR

Reverse transcription quantitative real-time PCR (RT-qPCR) reactions were performed in a CFX96TM Real-Time System, C1000TM Thermal Cycler from BIO-RAD using SYBR Green Universal Assay (BIO-RAD, 1725271). In each RT-qPCR well, 18 μL of mix were added, the mix contained 10 μl of SYBR Green 2X, 0,6 μl of a forward and reverse primer at 10 μM, and 7,4 μl of water. The primers had a final concentration of 0,3 μM/reaction. 2 μl containing 10 ng of cDNA were added to each well. Normalized values were calculated by dividing the mean expression value by a factor equal to the geometric mean of the normalization genes (*i*.*e*. beta-2 microglobulin *(B2M)*, transferrin receptor *(TFRC)* for MCF-7, SK-BR-3, MDA-MB-231, *B2M* and glyceraldehyde-3-phosphate dehydrogenase *(GAPDH)* for the Wood cells, and *B2M* for PB cells) and applying the ΔΔCt method.^3^ For a precise calculation, normalized values were obtained by dividing the expression value by a factor equal to the geometric mean of the normalization genes. Table 1 contains the complete list of primers used in this study.

### RNA-sequencing and data processing

For each condition, to ensure reliable statistical power, three independent replicates were prepared. To guarantee the extraction of enough total RNA from the samples lysed on day seven, five FN-silk networks were pulled together for each replicate. RNA integrity was determined by capillary electrophoresis on an Agilent 2100 Bioanalyzer using the Eukaryote Total RNA Nano assay, an RNA Integrity Number (RIN) major than 6 was observed for all samples. 200 ng of total RNA was used to generate stranded mRNA libraries with the mRNA prep ligation kit according to the manufacturer’s instructions (20040534, Illumina, USA). The concentration and quality of the cDNA libraries were assessed by Qubit (Life Technologies, USA) and TapeStation, respectively. Sequencing was performed with Illumina RNA-seq (Nextseq 2000, P2 flowcell, Paired End, dual index). For data processing, Illumina bcl2fastq (v2.20.0) was used for base calling and demultiplexing. Read quality was checked for each sample using FastQC (v0.11.8). Reads were aligned to Ensemble GRCh38/hg38 reference genome using STAR (v2.6. Id). Read summarization to gene counts was performed using featureCounts (vl .5.1). Normalized counts were generated using DESeq2 (vl .28.1). The principal component analysis (PCA) quality test results did not reveal any outliers. To identify the genes modulated by cultivation on FN-silk, we performed, for each cell line, two pairwise differential analyses comparing a) cells cultured in FN-silk network or 2D for seven days (*i*.*e*., FN-silk network vs 2D at day 7) and b) cells cultured in FN-silk network for seven days as opposed to the initial cell population harvested from the tissueculture treated flask (*i*.*e*., FN-silk network day 7 *vs* control flask at day 0). For both comparisons, we applied a conservative significance threshold of 5% to the corrected *p*-value, associated with a log2 fold-change ≥ |± 0,38|. A gene was considered as differentially expressed (DEG) by culture in 3D when significantly modulated in both comparisons.

### Statistical analysis

For the statistical analysis, 2-way ANOVA with interaction was used to analyze the data in GraphPad. Differences between groups were considered statistically significant when P-value < 0,05.

